# Introducing NOMAD: A Field-Deployable Workflow for Rapid, On-Site Microbiome Analysis of Raw Bovine Milk Using Full-Length 16S rRNA Sequencing

**DOI:** 10.1101/2025.09.29.679195

**Authors:** Valeria R. Parreira, Melinda Precious, Gurleen Taggar, Dharamdeo Singh, Lawrence Goodridge

**Author notes:** **Corresponding author:** Valeria R. Parreira.

## Abstract

Raw bovine milk hosts a diverse microbiota that profoundly influences dairy product quality, safety, and shelf-life. However, current surveillance methods are time-intensive and often lack the taxonomic resolution needed for effective risk mitigation. To address this, we developed NOMAD (Nanopore-based On-site Microbiome Analysis of Diversity), a field-deployable workflow for rapid, high-resolution characterization of the raw milk microbiome using full-length 16S rRNA gene sequencing via Oxford Nanopore Technologies. Milk samples collected from a commercial dairy operation were processed using eight DNA extraction protocols, with Method (incorporation of EDTA and TE buffer) emerging as the optimal approach for microbial richness and DNA yield. Sequencing was performed on a MinION Mk1B platform, and bioinformatic analyses revealed that a 4-hour run was sufficient to recover >90% of total community richness, with stable alpha and beta diversity metrics by this timepoint. The complete workflow, including DNA extraction, library preparation, sequencing, and analysis, was completed in 10.5 hours, enabling same-day microbiome profiling in farm-adjacent settings. Comparative analysis showed strong agreement with established milk microbiome studies, while full-length reads enhanced resolution of spoilage-associated taxa such as Pseudomonas spp. and Streptococcus spp. The NOMAD platform offers a powerful and practical tool for near real-time microbiological surveillance in the dairy industry, supporting proactive quality control and improved food safety outcomes.

## 1. INTRODUCTION

Raw bovine milk is a complex biological fluid that harbors diverse microbial communities influencing dairy product quality, safety, and shelf-life. Despite milk leaving the udder with relatively low microbial contamination, its microbiota is rapidly shaped by environmental exposures, milking equipment, storage conditions, and animal health (Martin et al., 2023). Certain bacterial groups, particularly psychrotrophic *Pseudomonas* spp. and spore-forming *Clostridium* spp. are of special concern due to their capacity to produce heat-stable lipases and proteases, leading to off-flavors, texture defects, and spoilage even after pasteurization (Martin et al., 2023; Murphy et al., 2016).

Traditionally, dairy microbiological surveillance has relied on culture-based methods and laboratory-bound assays such as standard plate counts (SPC) and flow cytometry, which are time-consuming and lack taxonomic resolution (Marri et al., 2020). These delays are suboptimal in the context of modern dairy supply chains that demand immediate corrective actions to maintain quality and compliance. The need for rapid, accurate, and portable microbial diagnostics is further underscored by recent studies that demonstrate significant seasonal and weather-driven shifts in milk microbiota, which can impact product performance and safety if undetected (Li et al., 2018).

To address these limitations, mobile genomics platforms, particularly the Oxford Nanopore Technologies (ONT) MinION, have emerged as promising tools for in situ analysis of the microbiome. When paired with portable lab equipment such as the Bento Lab or VolTRAX, these devices enable field-ready workflows that can sequence DNA and generate real-time results outside of laboratory environments (Bloemen et al., 2023; Chang et al., 2020). ONT-based mobile sequencing has been successfully deployed in remote settings, including pathogen surveillance in poultry, biodiversity assessments in marine ecosystems, and genome sequencing on the International Space Station, validating its resilience and flexibility (Bloemen et al., 2023; Chang et al., 2020).

A key advantage of ONT sequencing is its capacity for full-length 16S rRNA gene sequencing (∼1,500 bp), allowing for improved bacterial taxonomic resolution compared to conventional short-read (e.g., V3–V4) amplicon methods. This is especially critical in dairy microbiology, where differentiating between closely related spoilage organisms or commensals can inform targeted interventions (Bloemen et al., 2023). Long-read sequencing also facilitates a more robust linkage between genetic markers and functional traits, such as antimicrobial resistance genes or virulence factors, which are increasingly important in the context of One Health and food safety (Bloemen et al., 2023).

Recent studies have demonstrated the feasibility of portable sequencing workflows in identifying key dairy-relevant microbes, achieving turnaround times of less than 9 hours for DNA barcoding (Chang et al., 2020), and under 5 hours for *Campylobacter* detection in poultry using 16S rRNA amplicon sequencing (Bloemen et al., 2023). By comparison, traditional culture and qPCR assays may take 48–72 hours or longer, delaying risk mitigation actions at the farm or processing level.

In this study, we introduce NOMAD (Nanopore-based On-site Microbiome Analysis of Diversity), a novel mobile workflow tailored for the on-site, full-length 16S rRNA-based characterization of raw milk microbiomes. NOMAD is designed to operate in farm-adjacent environments using minimal infrastructure and delivers high-resolution taxonomic data in hours rather than days. By facilitating immediate microbial insight at the source, NOMAD offers a powerful tool for dairy producers, quality control managers, and food safety professionals aiming to detect spoilage risks, monitor hygiene practices, and ultimately enhance the microbiological quality of milk destined for processing.

## 2. MATERIAL AND METHODS

### 2.1 Sample collection and processing

Fresh raw bovine milk was collected in summer 2025 from the University of Guelph dairy farm. Bulk milk was sampled from the storage tank, transferred into sterile bottles, and transported to the laboratory on ice. In the laboratory, 45 mL aliquots were dispensed into 50 mL conical centrifuge tubes and stored at 4 °C for immediate processing or −80 °C for later use. Eight methods were evaluated in triplicate to optimize bacterial cell recovery, and workflows are summarized in Table 1.

**Table 1.** Overview of eight DNA extraction methods for raw bovine milk. Methods are compared based on storage conditions, sample volume, centrifugation speed, milk fraction processed, and use of bead beating. Each method was performed in triplicate.

| Method<sup>1</sup> | Sample storage | Milk Volume | Centrifugation speed<sup>2</sup> | Milk fraction extracted | Bead beating<sup>3</sup> |
| --- | --- | --- | --- | --- | --- |
| 1 | 4°C for 6 weeks | 45 ml | 8.000 x *g* | 250 mg fat | yes |
| 2 | -80°C | 45 ml | 8.000 x *g* | 250 mg fat | no |
| 3 | -80°C | 45 ml | 8.000 x *g* | whole pellet | no |
| 4 | -80°C | 45 ml | 8.000 x *g* | 250 mg fat | yes |
| 5 | -80°C | 7.2 ml | 8.000 x *g* | 250 mg fat + whole pellet | no |
| 6 | -80°C | 7.2 ml | 20.000 x *g* | 250 mg fat + whole pellet | no |
| 7 | -80°C | 45 ml | 8.000 x *g* | whole pellet | yes |
| 8 | -80°C | 45 ml | 8.000 x *g* | 250 mg fat + whole pellet | yes |

<sup>1</sup> Each extraction method was performed in triplicates.
<sup>2</sup> Centrifugation at 10 min, room temperature.
<sup>3</sup> Bead beating at 4800 rpm, for 1 min, repeated three times, with cooling on ice for 3 min between cycles.

Two sample processing protocols, based on the volume of milk tested, were evaluated: a 45 mL milk protocol (Methods 1, 2, 3, 4, 7, and 8) and a 7.2 mL milk protocol (Methods 5 and 6). Both approaches were designed to optimize bacterial recovery while accounting for sample volume and downstream processing requirements.

45 mL milk protocol (Methods 1, 2, 3, 4, 7, and 8). Each 45 mL milk sample was supplemented with 3 mL of 500 mM EDTA (pH 8.0) and 2 mL of TE buffer (pH 7.6), then mixed by inversion to disrupt casein micelles. Samples were centrifuged at 8.000 × g for 10 min at room temperature. For fat samples, 250 mg of solidified fat was transferred to a processing tube using a sterile 10 µL disposable loop. For pellet-based methods, all whey was removed, and the pellet was resuspended in 200 µL of lysis buffer (PM1, AllPrep PowerViral DNA/RNA Kit, Qiagen) before transfer to a processing tube containing an additional 700 µL of PM1 buffer.

7.2 mL milk protocol (Methods 5 and 6). Four 1.8 mL aliquots were taken from a 45 mL tube (after EDTA and TE addition) and centrifuged in 2 mL tubes for 10 min at room temperature. Centrifugation was performed at either 8000 × g or 20,000 × g, depending on the method (see Table 1). Whey fractions were removed, and the pellets and fat from all four tubes were combined and suspended in a single volume of 900 µL of PM1 buffer.

### 2.2 Genomic DNA Extraction

Total genomic DNA was extracted using the AllPrep PowerViral DNA/RNA Kit (Qiagen, Hilden, Germany) following the manufacturer’s protocol, with specific modifications to optimize recovery. All centrifugation steps were conducted at 8,000 × g using the BentoLab portable centrifuge (BentoLab, UK). The final column-drying spin was extended to 5 minutes to ensure complete removal of residual ethanol prior to elution.

For protocols involving mechanical lysis, 900 µL of lysis buffer PM1 was added to each sample in 2 mL screw-cap tubes containing 2 g of 0.1 mm zirconia/silica beads (Biospec Products Inc., USA). Samples were incubated at 80°C for 10 minutes, briefly cooled on ice (2 minutes), and subjected to bead beating at 4,800 rpm for 1 minute, repeated three times with 3-minute cooling intervals on ice between each cycle. For methods without bead beating, samples were incubated at 80°C for 10 minutes post-resuspension in PM1 buffer before proceeding. A total of 700 µL of cleared lysate was used for DNA purification. The protocol was further modified by incorporating an additional 400 µL wash step with both PM4 and PM5 buffers to improve DNA purity. Elution was performed using 85 µL of nuclease-free water.

DNA quality and concentration were evaluated for all samples. Purity was assessed spectrophotometrically (260/280 and 260/230 absorbance ratios) using the QIAxpert system (Qiagen), and DNA concentration was quantified using the Qubit dsDNA High Sensitivity Assay Kit with the Qubit 4 Fluorometer (Invitrogen, USA), according to the manufacturer’s instructions.

### 2.3 16S rRNA Gene Amplicon Sequencing

The entire region of the bacterial 16S rRNA gene was amplified from extracted DNA using the 16S Barcoding Kit 24 V14 (SQK-16S114.24) (Oxford Nanopore Technologies, UK) following the manufacturer’s recommendations. PCR was performed for 30 cycles with an annealing temperature of 55°C, using 50 ng of template DNA per reaction. Negative controls (nuclease-free water) were included to monitor potential contamination.

Amplicons were purified using AMPure XP beads (Beckman Coulter, Canada) and quantified with the Qubit 4 Fluorometer using the dsDNA HS Assay Kit. Libraries were normalized to a final concentration of 50–100 fmol before sequencing. Sequencing was carried out on a MinION Mk1B platform using FLO-MIN114 (R10.4.1) flow cells, with a runtime of 72 hours.

### 2.4 Determination of Minimum Sequencing Run Time for the Oxford Nanopore 16S Workflow

To assess the minimal sequencing duration required for robust taxonomic profiling, Bioinformatic analyses were conducted at 2-hour intervals up to 24 hours and at 72 hours, with sequence files from each interval compared to the complete 72-hour dataset. Read subsets were generated based on start time metadata from the sequencing summary file provided by ONT MinKNOW. Taxonomic profiles at each time point were compared to the full 72-hour dataset to evaluate completeness and consistency.

### 2.5 Bioinformatic Processing and Taxonomic Assignment

Real-time basecalling was performed using the MinKNOW GUI (v24.11.8) with the Dorado basecaller (model: dna_r10.4.1_e8.2_400bps_hac, version 4.3.0). Quality control was implemented using NanoFilt, with reads filtered for an average Phred score ≥ 8, minimum read length of 100 bp, and trimming of low-quality terminal bases.

Taxonomic classification of high-quality reads was performed with Kraken2 (Wood et al., 2019) against the Kraken2 reference database. Species-level relative abundances were refined using Bracken v2.9, configured with a 100 bp k-mer size to match the corresponding Bracken database. An abundance threshold of 10% was applied to reduce noise from rare taxa.

### 2.6 Statistical Analysis

Statistical analyses were conducted to evaluate the reproducibility and consistency of microbial community profiles across DNA extraction methods and sequencing durations (Supplemental methods_merged.bracken_S.counts+percent.csv). Replicates were averaged per method (supplemental method_abundance_raw.csv), and a 1% relative abundance threshold was applied to filter out low-abundance taxa (supplemental method_abundance_filtered.csv). Alpha diversity was quantified using species richness, Shannon index, and Simpson index Beta diversity was assessed using both Bray–Curtis dissimilarity (abundance-weighted) and Jaccard index (presence/absence).

Ordination was performed via Principal Coordinates Analysis (PCoA), and clustering analysis was conducted using hierarchical clustering. Differences in community composition across methods were evaluated using PERMANOVA (adonis2 function), based on both Bray–Curtis and Jaccard distance matrices. Significant, pairwise PERMANOVA was conducted to identify method-specific differences.

The effect of sequencing duration on community profiling was assessed using similar methods (supplementary_all_times_merged.bracken_S.counts+percent.csv and 72hmerged.bracken_S.counts+percent.csv). Alpha diversity metrics were calculated for each time point using species richness, Shannon’s index, and Simpson’s index, after applying a 1% relative abundance threshold. Beta diversity was assessed using Bray–Curtis and Jaccard indices. Spearman rank correlations were calculated between profiles at each time point and the 72-hour endpoint to quantify temporal convergence. All statistical analyses and data visualizations were conducted in RStudio using the vegan package (v2.7-1) and ggplot2 (v3.5.2).

## 3. RESULTS

### 3.1 Comparison of Extraction Methods for Microbial Community Recovery

Eight DNA extraction methods were evaluated to optimize bacterial cell recovery from raw milk, with triplicate preparations performed for each method. Sequencing was performed using the Oxford Nanopore Technologies SQK-16S114 kit with a 72-hour runtime. Alpha and beta diversity analyses were used to assess the taxonomic richness and structural characteristics of the microbial communities recovered by each method. Tables 1–4, Figure 1, and Supplementary Figures 1–3 compare eight DNA extraction methods, highlighting differences in microbial recovery, community richness, and taxonomic composition.

**Figure 1.**
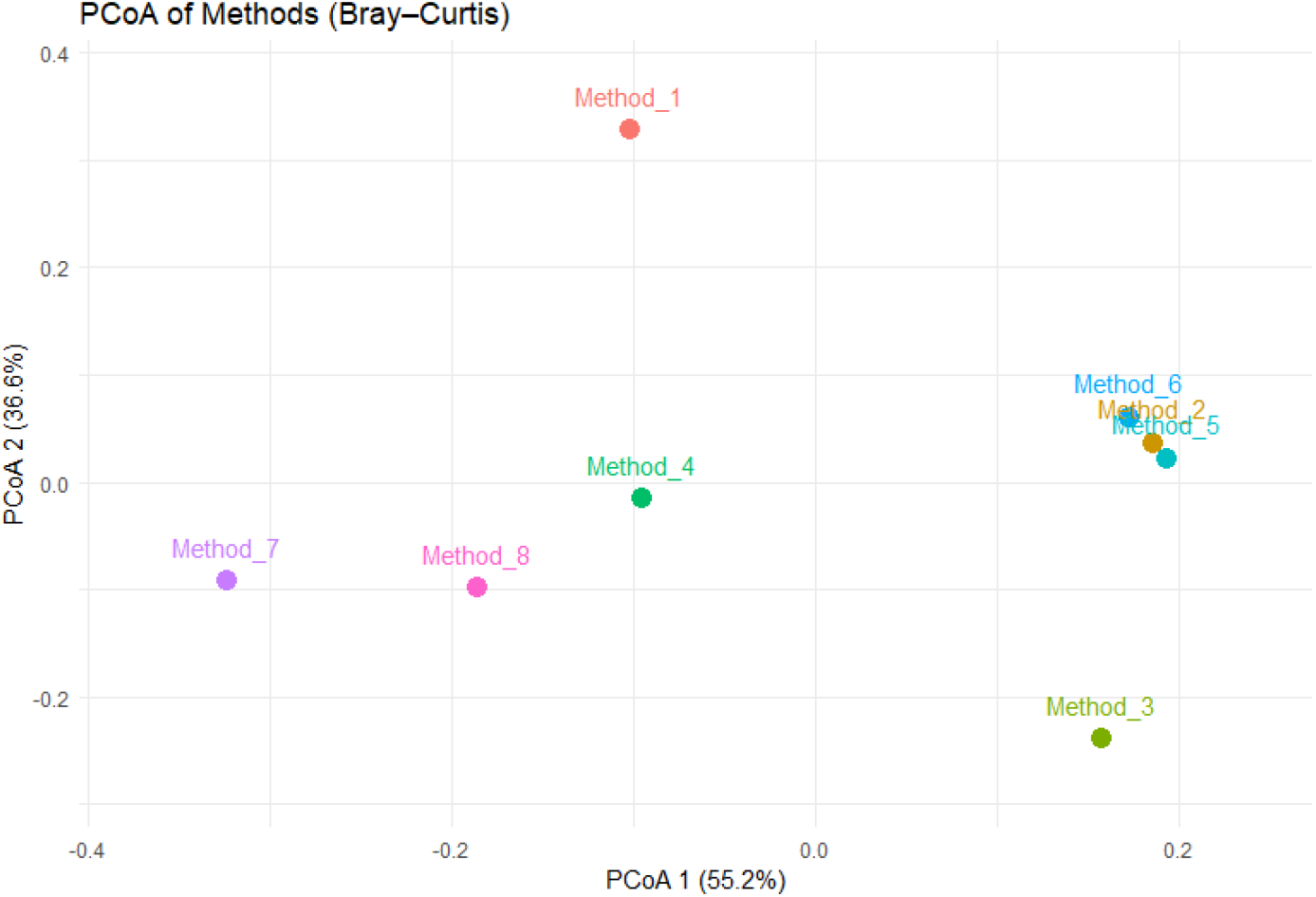
Principal Coordinates Analysis (PCoA) plot of Bray–Curtis distances showing clustering of microbial communities obtained with the eight DNA extraction methods.

Significant variation in species richness and diversity was observed across the eight methods. Richness ranged from 90 species in Method 1 to 162 species in Method 3, while Shannon and Simpson indices highlighted differences in community evenness and dominance (Table 2). Methods 3, 4, 6, 7, and 8 yielded the highest Shannon values (2.8–2.93), indicating balanced community structures. Method 1 consistently performed the poorest, with the lowest richness (90 species) and Shannon index (2.40). Simpson indices reinforced these findings, with the most even communities (Simpson > 0.90) observed in Methods 3, 4, 5, and 8. These results suggest that while no single method was optimal across all metrics, Methods 3, 4, 5, and 8 emerged as the strongest performers in terms of taxonomic breadth and balance.

**Table 2.** Alpha diversity indices (species richness, Shannon, Simpson) of microbial communities recovered using the eight DNA extraction methods.

| Method | Richness | Shannon | Simpson |
| --- | --- | --- | --- |
| Method_1 | 90 | 2.395054 | 0.858434 |
| Method_2 | 121 | 2.725716 | 0.892521 |
| Method_3 | 162 | 2.916221 | 0.90965 |
| Method_4 | 127 | 2.900013 | 0.907883 |
| Method_5 | 145 | 2.887992 | 0.905039 |
| Method_6 | 139 | 2.844112 | 0.898064 |
| Method_7 | 132 | 2.804544 | 0.883183 |
| Method_8 | 131 | 2.93204 | 0.902654 |

Beta diversity analysis revealed method-dependent shifts in microbial community composition. Both Bray-Curtis and Jaccard dissimilarity metrics detected statistically significant differences across methods (Bray-Curtis PERMANOVA: R^2^ = 0.59, p = 0.001; Jaccard PERMANOVA: R^2^ = 0.63, p = 0.001). Principal Coordinates Analysis (PCoA) of Bray-Curtis distances (Figure 1) captured 92% of total variation on the first two axes, emphasizing strong method-driven differences. Method 1 was clearly separated from all others, while Methods 2, 5, and 6 formed a tight cluster, indicating similar taxonomic profiles. Method 3 resolved independently, suggesting distinct recovery biases. Methods 7 and 8 clustered together, and Method 4 occupied an intermediate position between the central and peripheral clusters.

Bray–Curtis dissimilarity values (Table 3) indicated that Method 1 was the most divergent from other methods, while Methods 4, 6, and 8 exhibited the highest compositional similarity. In contrast, Jaccard distances (Table 4) highlighted greater variability in shared taxa, particularly between Method 3 and the remaining methods. These distance-based differences were consistent with pairwise PERMANOVA findings, which highlighted statistically significant differences between Method 1 and Method 4 (p = 0.029) for Jaccard and (p=0.031) for Bray Curtis, and between Method 3 and Method 4 (p = 0.039) for Jaccard and (p=0.03) for Bray Curtis, further underscoring the taxonomic specificity imparted by extraction method (Supplementary Methods_Pairwise_PERMANOVA_BrayCurtis.csv and Methods_Pairwise_PERMANOVA_Jaccard.csv).

**Table 3.**
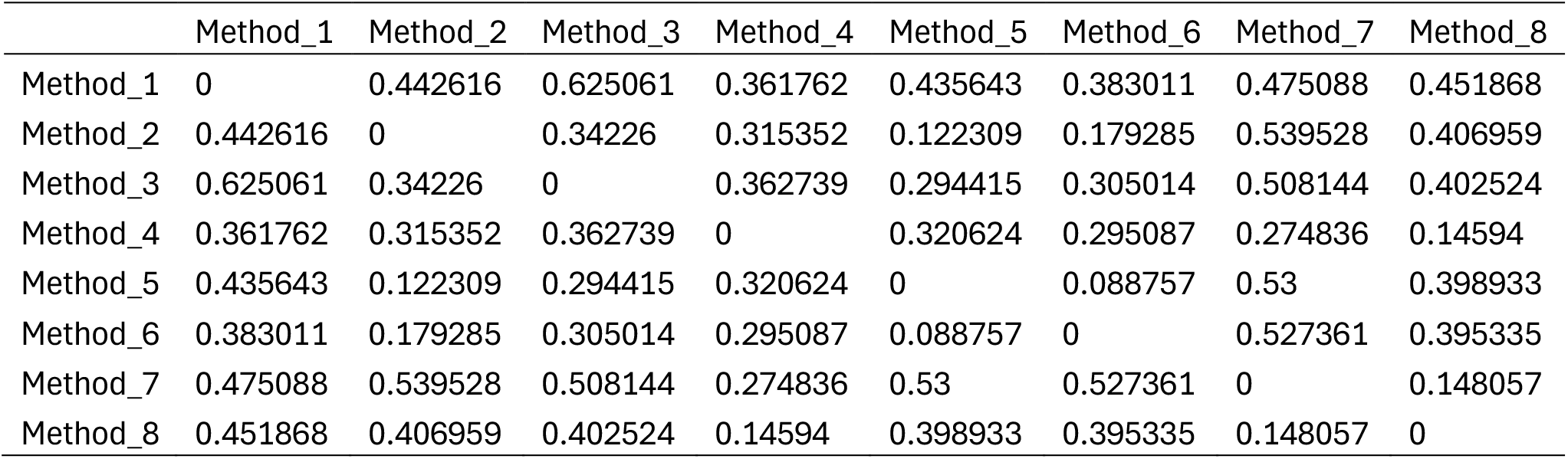
Pairwise Bray–Curtis dissimilarity values comparing microbial community composition across the eight DNA extraction methods.

|  | Method_1 | Method_2 | Method_3 | Method_4 | Method_5 | Method_6 | Method_7 | Method_8 |
| --- | --- | --- | --- | --- | --- | --- | --- | --- |
| Method_1 | 0 | 0.442616 | 0.625061 | 0.361762 | 0.435643 | 0.383011 | 0.475088 | 0.451868 |
| Method_2 | 0.442616 | 0 | 0.34226 | 0.315352 | 0.122309 | 0.179285 | 0.539528 | 0.406959 |
| Method_3 | 0.625061 | 0.34226 | 0 | 0.362739 | 0.294415 | 0.305014 | 0.508144 | 0.402524 |
| Method_4 | 0.361762 | 0.315352 | 0.362739 | 0 | 0.320624 | 0.295087 | 0.274836 | 0.14594 |
| Method_5 | 0.435643 | 0.122309 | 0.294415 | 0.320624 | 0 | 0.088757 | 0.53 | 0.398933 |
| Method_6 | 0.383011 | 0.179285 | 0.305014 | 0.295087 | 0.088757 | 0 | 0.527361 | 0.395335 |
| Method_7 | 0.475088 | 0.539528 | 0.508144 | 0.274836 | 0.53 | 0.527361 | 0 | 0.148057 |
| Method_8 | 0.451868 | 0.406959 | 0.402524 | 0.14594 | 0.398933 | 0.395335 | 0.148057 | 0 |

**Table 4.** Pairwise Jaccard distances showing presence/absence-based differences in microbial community membership across extraction methods.

|  | Method_1 | Method_2 | Method_3 | Method_4 | Method_5 | Method_6 | Method_7 | Method_8 |
| --- | --- | --- | --- | --- | --- | --- | --- | --- |
| Method_1 | 0 | 0.492857 | 0.673684 | 0.553333 | 0.575758 | 0.532051 | 0.6125 | 0.592357 |
| Method_2 | 0.492857 | 0 | 0.453552 | 0.312925 | 0.25 | 0.243243 | 0.502959 | 0.374194 |
| Method_3 | 0.673684 | 0.453552 | 0 | 0.420765 | 0.433673 | 0.407407 | 0.444444 | 0.344633 |
| Method_4 | 0.553333 | 0.312925 | 0.420765 | 0 | 0.371257 | 0.294872 | 0.329032 | 0.183099 |
| Method_5 | 0.575758 | 0.25 | 0.433673 | 0.371257 | 0 | 0.213836 | 0.510753 | 0.385965 |
| Method_6 | 0.532051 | 0.243243 | 0.407407 | 0.294872 | 0.213836 | 0 | 0.502762 | 0.363636 |
| Method_7 | 0.6125 | 0.502959 | 0.444444 | 0.329032 | 0.510753 | 0.502762 | 0 | 0.281046 |
| Method_8 | 0.592357 | 0.374194 | 0.344633 | 0.183099 | 0.385965 | 0.363636 | 0.281046 | 0 |

Among the eight methods, Method 4 stood out as a balanced performer, achieving high richness (127 species), a Shannon index of 2.90, and a Simpson index of 0.91. In ordination and clustering analyses, it remained compositionally distinct from outliers like Method 1 and Method 3, while aligning more closely with mid-performing methods (2, 5, and 6).

### 3.2 Sequencing Time Optimization for Field Applications

To determine the minimum ONT sequencing time needed to capture a stable milk microbiota profile, one raw milk sample (processed using Method 4) was sequenced in triplicate and analyzed at 13 different timepoints from 2 to 72 hours. Tables 5–7 and Figures 2–3 evaluate sequencing duration, showing that stable microbial diversity profiles are achieved by 4 hours, closely matching the 72-hour reference dataset. For example, as shown in table 5, species richness increased rapidly in the early timepoints, with 105 species (85% of total richness) detected at 2 hours. By 4 hours, 115 species (93%) were identified. Shannon and Simpson indices stabilized rapidly, with minimal changes between 4 hours and the 72-hour endpoint, suggesting that the majority of ecological structure was captured early in the run (Figure 2).

**Table 5.** Alpha diversity indices (species richness, Shannon, Simpson) of microbial communities in raw milk sequenced at multiple timepoints (2–72 h), benchmarked against the 72 h dataset.

| time_hr | richness | shannon | simpson |
| --- | --- | --- | --- |
| 2 | 105 | 2.123712 | 0.786556 |
| 4 | 115 | 2.13734 | 0.788131 |
| 6 | 117 | 2.140909 | 0.788137 |
| 8 | 120 | 2.144192 | 0.788023 |
| 10 | 123 | 2.147997 | 0.787834 |
| 12 | 121 | 2.144715 | 0.787487 |
| 14 | 124 | 2.146464 | 0.787302 |
| 16 | 125 | 2.145857 | 0.786966 |
| 18 | 126 | 2.145687 | 0.786616 |
| 20 | 126 | 2.145213 | 0.786192 |
| 22 | 127 | 2.14721 | 0.785845 |
| 24 | 124 | 2.143842 | 0.785316 |
| 72 | 124 | 2.124551 | 0.776401 |

**Table 6.** Bray–Curtis dissimilarity and Jaccard distance values comparing microbial community composition at each sequencing timepoint to the 72 h dataset.

| time_hr | bray_vs_72 | jaccard_vs_72 |
| --- | --- | --- |
| 2 | 0.038156 | 0.238461538 |
| 4 | 0.037688 | 0.175572519 |
| 6 | 0.036905 | 0.160305344 |
| 8 | 0.036231 | 0.151515152 |
| 10 | 0.03508 | 0.128787879 |
| 12 | 0.034583 | 0.143939394 |
| 14 | 0.033888 | 0.121212121 |
| 16 | 0.032693 | 0.127819549 |
| 18 | 0.031498 | 0.106060606 |
| 20 | 0.030064 | 0.106060606 |
| 22 | 0.028649 | 0.098484848 |
| 24 | 0.027508 | 0.07751938 |
| 72 | 0 | 0 |

**Figure 2.**
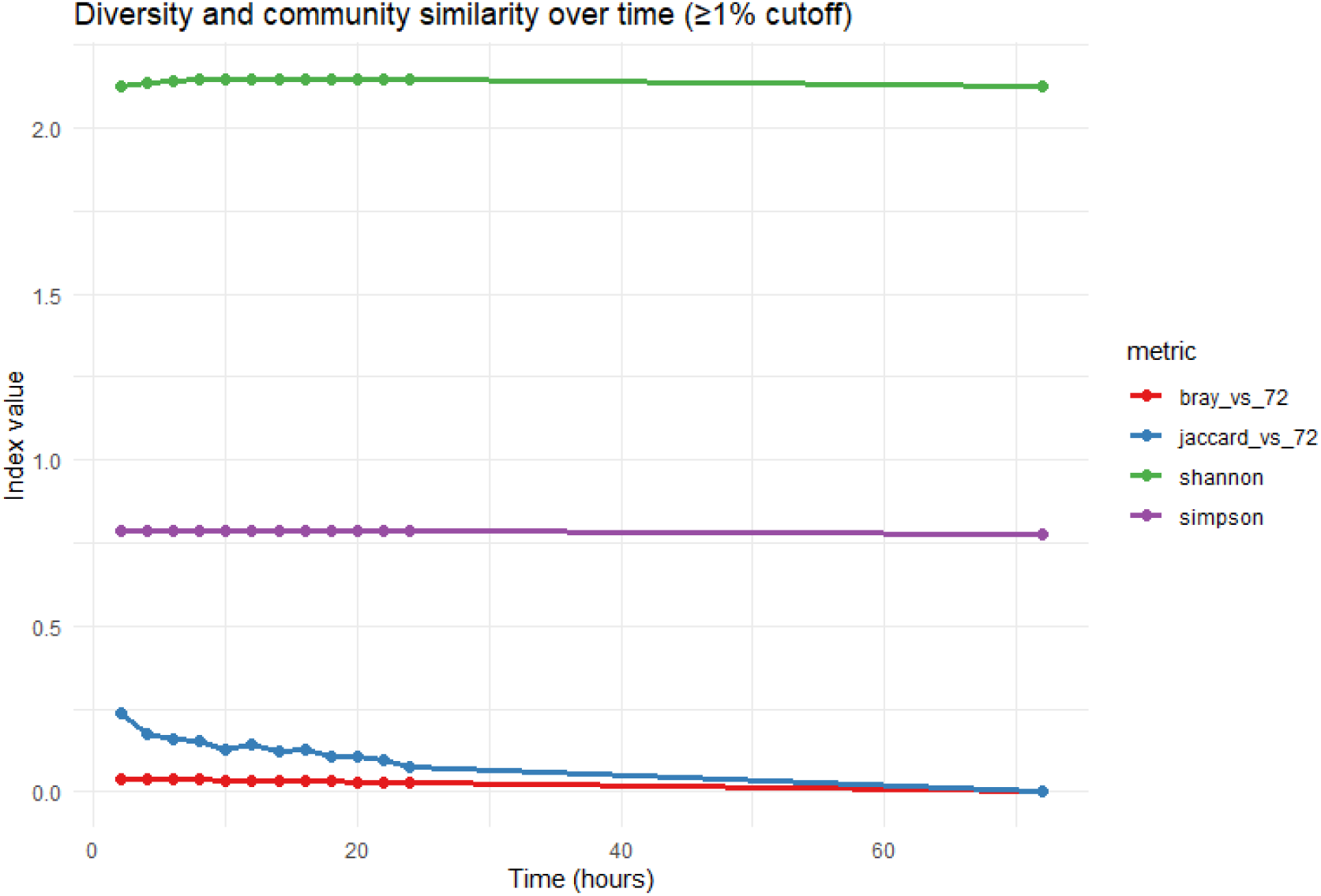
Alpha and beta diversity metrics (species richness, Shannon, Simpson, Bray– Curtis, Jaccard) across sequencing timepoints (2–72 h).

Species turnover analysis revealed 19 net additions between 2 and 72 hours. Late-appearing taxa included *Staphylococcus* sp. T93, *Bacteroides thetaiotamicron*, and *Vesimonas coprocola*. These were mostly low- to mid-abundance species that did not substantially alter the core community structure. Early-appearing but later-absent taxa, such as *Pseudomonas syringae* and *Pseudomonas* sp. FP435, which were likely transient organisms, diminished over time as community stabilization occurred. By 12–16 hours, taxonomic additions plateaued, and the core profile became stable (Supplementary species_gain_loss_all_vs_72h.csv).

Bray-Curtis dissimilarity values between each timepoint and the 72-hour endpoint showed a progressive convergence (Figure 3), dropping from 0.038 at 2 h to <0.03 by 20 h. Jaccard distances (presence/absence) followed a similar trend (Figure 3), declining to 0.078 by 24 h. Importantly, PERMANOVA on both Bray-Curtis and Jaccard matrices did not detect significant compositional differences across timepoints (p = 0.999 and p = 0.998, respectively), confirming minimal structural divergence. Spearman correlation of species rank abundance (Table 7) reached 0.87 at 4 hours and exceeded 0.94 by 22 hours, indicating strong concordance with the final 72-hour community.

**Table 7.** Spearman rank correlation coefficients comparing species abundance profiles at each sequencing timepoint (2–72 h) with the 72 h dataset.

| time_hr | spearman_vs_72 |
| --- | --- |
| 2 | 0.80779 |
| 4 | 0.866073 |
| 6 | 0.862911 |
| 8 | 0.874061 |
| 10 | 0.891201 |
| 12 | 0.893728 |
| 14 | 0.895695 |
| 16 | 0.911726 |
| 18 | 0.913109 |
| 20 | 0.92273 |
| 22 | 0.94287 |
| 24 | 0.945785 |
| 72 | 1 |

**Figure 3.**
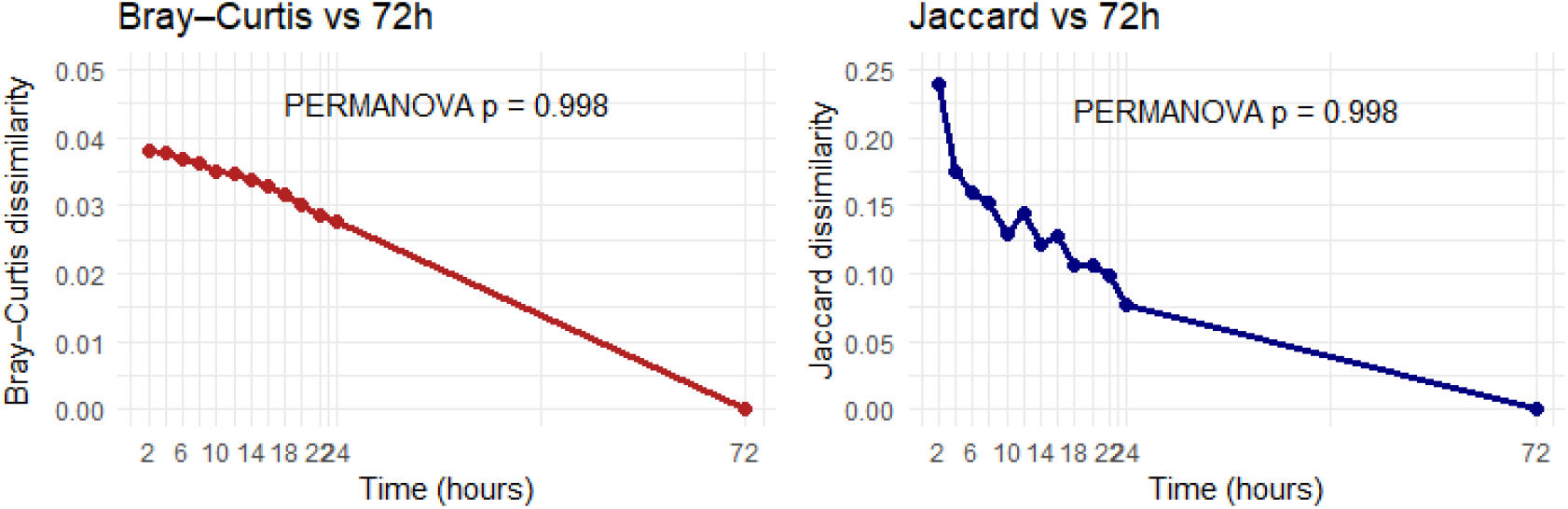
Statistical comparison of beta diversity across sequencing timepoints.

The top 20 most abundant taxa remained stable throughout the sequencing timeline (Supplemental top20_summary_overlap.csv). Core milk-associated species such as *Streptococcus agalactiae, Leuconostoc lactis*, and *Lactobacillus delbrueckii* were consistently dominant. From 4 hours onward, the identity and relative abundance of dominant taxa closely resembled the endpoint, indicating that deeper sequencing mainly contributed to the detection of low-abundance organisms without altering the ecological core.

## 4. DISCUSSION

The microbiota of raw milk represents a critical yet often under-monitored aspect of dairy production, directly influencing product quality, safety, shelf-life, and consumer health. In this study, we demonstrated the feasibility and utility of NOMAD—a mobile, full-length 16S rRNA sequencing workflow optimized for rapid, on-site microbiome analysis of raw bovine milk. By deploying the Oxford Nanopore MinION platform in conjunction with a streamlined sample preparation protocol (Method 4) and a 4-hour sequencing run, we achieved high-resolution microbial profiling within a total workflow of 10.5 hours. This approach aligns with the growing need for decentralized and real-time diagnostics in dairy supply chains. When compared to traditional lab-based and culture-dependent approaches, NOMAD offers not only speed and portability but also deeper taxonomic resolution, enabling producers and food safety professionals to detect and respond to microbial shifts at the source. The findings from this study, when compared with established literature, provide essential insights into the microbial ecology of raw milk and reinforce the value of rapid, field-deployable sequencing in modern dairy microbiology.

The microbial communities identified in raw bovine milk using the NOMAD workflow were broadly consistent with prior studies, yet demonstrated distinctive compositional features that likely reflect local environmental and methodological differences. As reported in earlier work (Yuan et al., 2022; Li et al., 2018), the predominant phyla in raw milk included Proteobacteria, Firmicutes, and Bacteroidota, with *Pseudomonas, Acinetobacter, Streptococcus*, and *Lactobacillus* among the most abundant genera. These taxa represent a mix of spoilage-associated organisms and beneficial commensals or fermenters, which are central to dairy quality outcomes.

However, our samples collected during the summer months revealed a notably higher relative abundance of Pseudomonas spp., consistent with the proliferation of psychrotrophs under cool storage conditions. This pattern was more pronounced than in studies conducted in cooler climates or under different housing systems (Quigley et al., 2013). At the same time, the presence of genera such as *Lactococcus* and *Leuconostoc i*n our samples suggests a microbial profile that may support early fermentation or reflect mild environmental cross-contamination. These findings highlight the dynamic and context-sensitive nature of the raw milk microbiota, which is influenced by seasonal factors, hygiene practices, storage temperatures, and feed management. Methodological differences likely contributed to the observed taxonomic profiles. While earlier studies typically employed short-read sequencing targeting the V3–V4 regions of the 16S rRNA gene, the use of full-length 16S sequencing via nanopore technology in the present study enhanced taxonomic resolution, particularly for differentiating closely related spoilage or pathogenic species. This distinction is crucial in dairy quality management, where slight taxonomic differences can have significant functional consequences.

Our adoption of full-length 16S rRNA gene sequencing using the Oxford Nanopore MinION platform enabled rapid, in-depth characterization of the raw milk microbiota at near-species level. This long-read approach provides distinct advantages over short-read methods, particularly in terms of resolving microbial diversity in complex matrices like milk, where spoilage organisms and beneficial microbes may co-occur within the same genus. For instance, the capacity to distinguish between *Pseudomonas fluorescens* and *Pseudomonas fragi* offers meaningful insight into spoilage potential and the likely enzymatic activity within a batch of milk. Compared to culture-based techniques and qPCR assays which typically require 48 to 72 hours and are constrained to specific target organisms NOMAD delivered actionable taxonomic profiles within five hours of sequencing. These findings are consistent with recent literature demonstrating the practicality and accuracy of nanopore-based microbial surveillance in on-site agricultural and environmental settings (Bloemen et al., 2023; Chang et al., 2020). Furthermore, the integration of real-time data analysis platforms such as EPI2ME allowed sequencing results to be interpreted without the need for extensive bioinformatics infrastructure, thus facilitating deployment outside conventional laboratory environments.

Sample preparation method 4 was selected for the final NOMAD workflow following comparative evaluation of eight sample preparation protocols. This method, which utilized a 45 mL milk input volume supplemented with EDTA and TE buffer, consistently yielded the highest DNA concentrations and microbial diversity. The chelating action of EDTA was instrumental in disrupting casein micelles and releasing tightly bound bacterial cells, thereby enhancing recovery from a protein-rich matrix.

The sequencing data revealed that a four-hour sequencing period is sufficient for achieving taxonomic depth while maintaining throughput and field feasibility. Longer sequencing runs offered only marginal improvements in classification depth but introduced diminishing returns in terms of time and power consumption, which are key constraints in on-site applications. Including DNA extraction, library preparation, sequencing, and analysis, the whole workflow was completed in approximately 10.5 hours, supporting same-day diagnostics and quality decision-making.

The field-deployable nature of the NOMAD pipeline offers a practical solution to microbiome surveillance in dairy environments where traditional laboratory access is limited or delayed. Equipped with the Bento Lab for molecular processing and the MinION for sequencing, the workflow requires only minimal infrastructure and can be operated in mobile labs or farm-adjacent settings. This portability enables producers to monitor microbial dynamics in real time and take immediate action to mitigate contamination risks. Application scenarios include routine profiling of bulk tank milk to assess hygiene status, early detection of psychrotrophic spoilage organisms prior to shipment, and longitudinal surveillance to trace contamination sources or evaluate the impact of environmental changes such as bedding type, feed composition, or seasonal shifts. Furthermore, integration with digital dashboards and cloud-based analytics could support centralized quality assurance systems, allowing dairy cooperatives and processors to optimize supply chain management based on microbial risk patterns.

By decentralizing access to high-resolution microbiome data, NOMAD presents a transformative tool for dairy producers and food safety professionals, bridging the gap between molecular diagnostics and practical on-farm decision-making. As the cost and accessibility of sequencing technologies continue to improve, such workflows are poised to become integral components of modern dairy management systems.

## 5. DATA AVAILABILITY

All raw sequencing reads generated in this study and processed taxonomic classification tables, species-level relative abundance matrices, and replicate-level alpha and beta diversity indices are available in Zenodo at DOI: 10.5281/zenodo.17221533 for the fresh milk timepoint comparison and DOI: 10.5281/zenodo.17222465 for the methods comparison. Supplementary figures and tables are included with the manuscript as supplementary files and are available in Zenodo at DOI: 10.5281/zenodo.17223101.

All analysis scripts used for diversity and ordination analyses were written in R (v4.3.0) and are available in the GitHub repository: https://github.com/singhdharamdeods/milk_analysis_methods_scripts and https://github.com/singhdharamdeods/python_scripts_milk_analysis/blob/main/Timepoints_Milk.R.

## 6. ACKNOWLEDGMENTS

The authors gratefully acknowledge support from the Canada Research Chairs Program, which provided funding for this work.

## Supplementary Figures

**Supplemental Figure 1.**
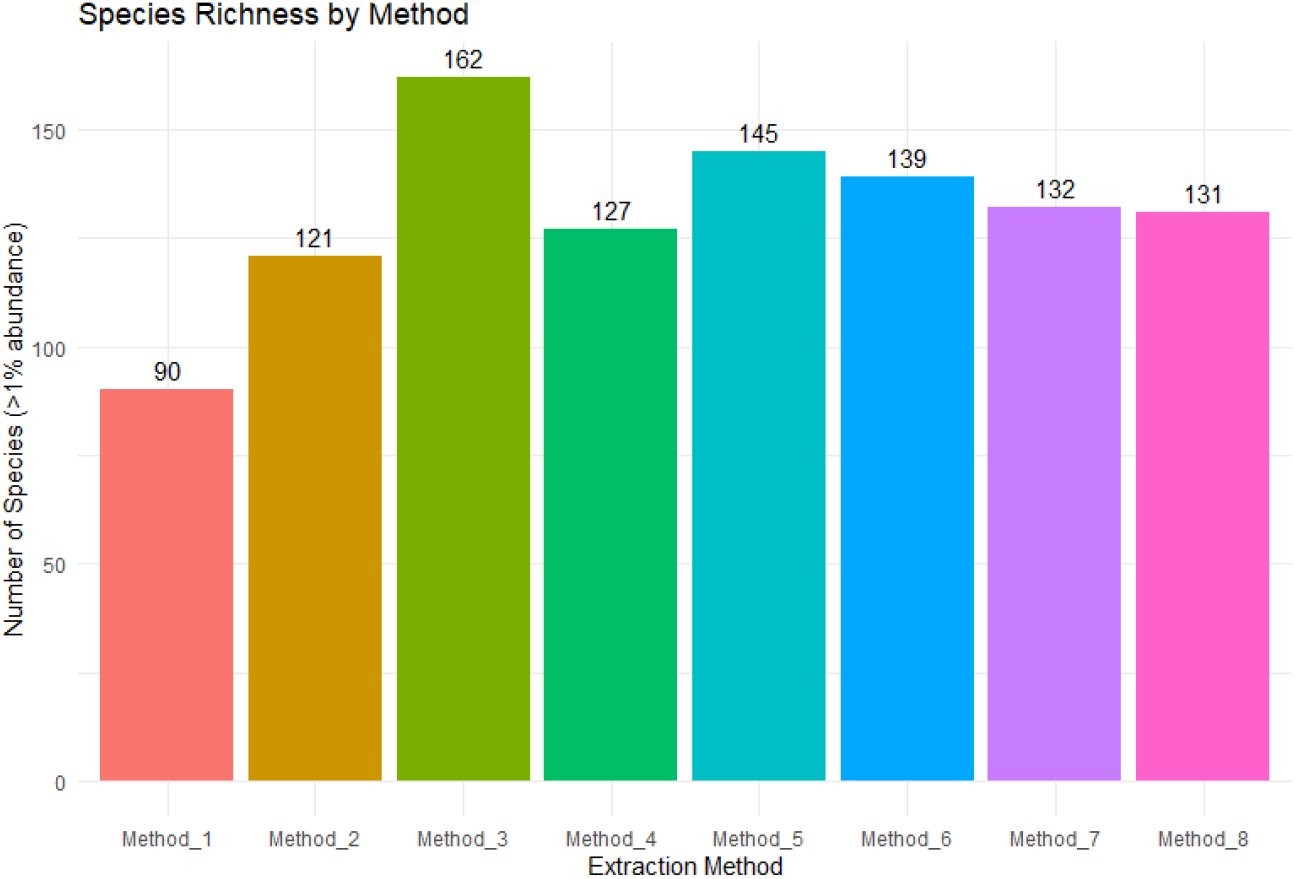
Species richness across the eight DNA extraction methods for raw bovine milk.

**Supplemental Figure 2.**
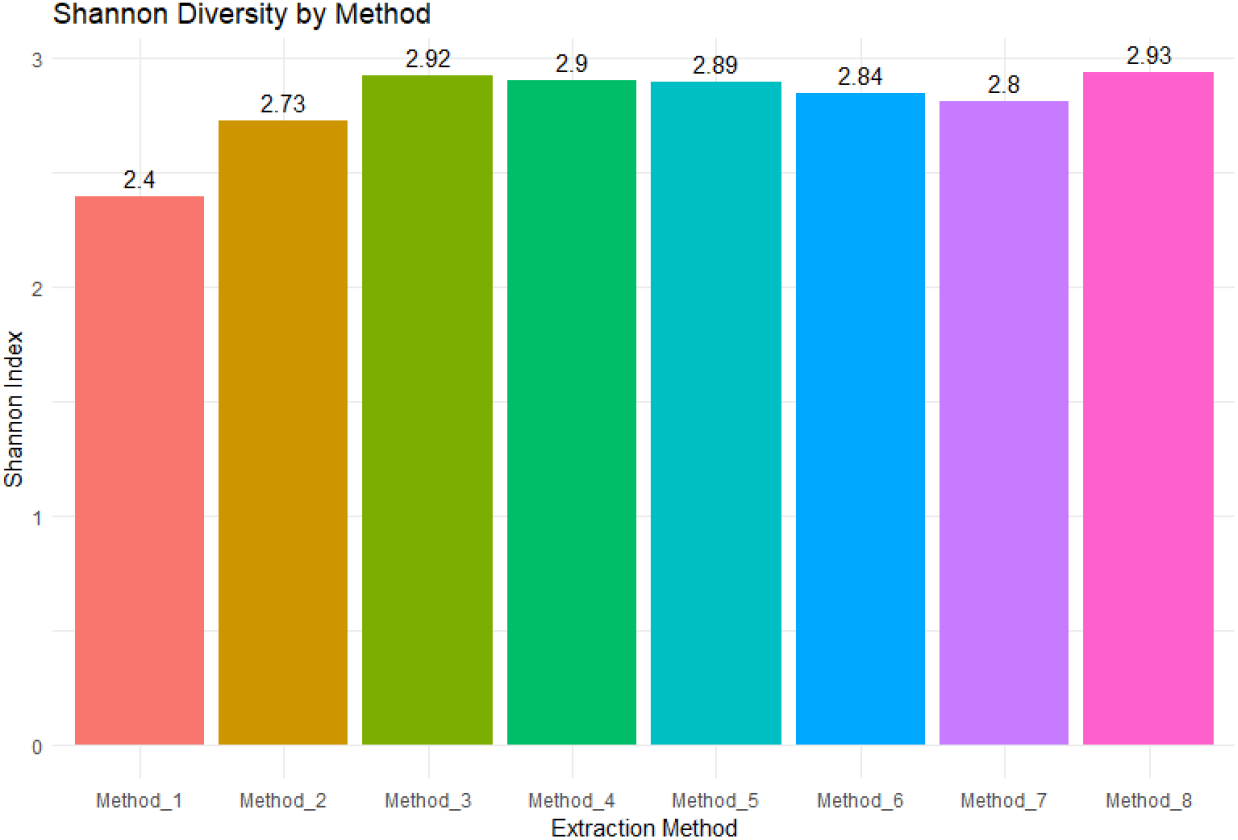
Shannon diversity indices across the eight DNA extraction methods.

**Supplemental Figure 3.**
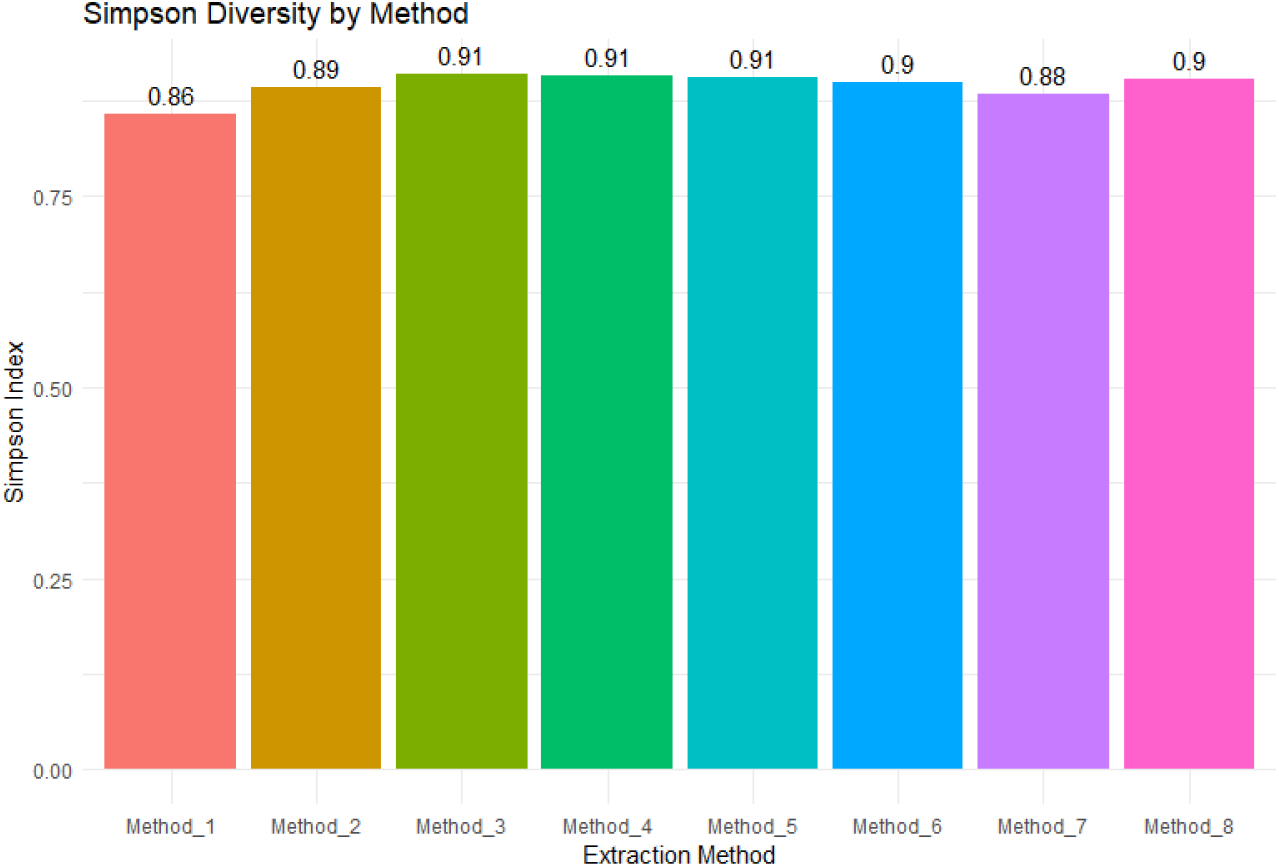
Simpson diversity indices across the eight DNA extraction methods.

